# Enhancing single-cell proteomics through tailored Data-Independent Acquisition and micropillar array-based chromatography

**DOI:** 10.1101/2022.11.29.518366

**Authors:** Valdemaras Petrosius, Pedro Aragon-Fernandez, Nil Üresin, Teeradon Phlairaharn, Benjamin Furtwängler, Jeff op de Beeck, Simon Francis Thomsen, Ulrich auf dem Keller, Bo T. Porse, Erwin M. Schoof

## Abstract

Single-cell resolution analysis of complex biological tissues is fundamental to capture cell-state heterogeneity and distinct cellular signaling patterns that remain obscured with population-based techniques. The limited amount of material encapsulated in a single cell however, raises significant technical challenges to molecular profiling. Due to extensive optimization efforts, mass spectrometry-based single-cell proteomics (scp-MS) has emerged as a powerful tool to facilitate proteome profiling from ultra-low amounts of input, although further development is needed to realize its full potential. To this end, we carried out comprehensive analysis of orbitrap-based data independent acquisition (DIA) for limited material proteomics. Notably, we found a fundamental difference between optimal DIA methods for high- and low-load samples. We further improved our low-input DIA method by relying on high-resolution MS1 quantification, thus more efficiently utilizing available mass analyzer time. With our ultra-low input tailored DIA method, we were able to accommodate long injection times and high resolution, while keeping the scan cycle time low enough to ensure robust quantification. Finally, we establish a complete experimental scp-MS workflow, combining DIA with accessible single-cell sample preparation and the latest chromatographic and computational advances and showcase our developments by profiling real single cells.

## Introduction

Analytical techniques with single-cell resolution are becoming indispensable tools to study complex biological systems. Although invaluable, the aggregated view obtained by bulk cell population experiments is not sufficient to achieve fundamental understanding of human development and disease. The means to interrogate the first two aspects of the central dogma of biology (DNA-RNA-Protein) are well established and have been widely adopted, but the study of proteomes by liquid chromatography coupled mass spectrometry (LC-MS) at single-cell resolution is just entering the biological application phase^1^. It is estimated that a single mammalian cell contains 50-450pg of protein^2^, posing significant challenges to protein identification and quantification. However, these challenges are to a large extent being mitigated by advances in different aspects of LC-MS-based proteomics ^3–13^.

Pioneering studies could quantify hundreds of proteins from a single cell^9,13^. These reports marked an important milestone for mass-spectrometry based single-cell proteomics (scp-MS), however analysis required long chromatographic gradients, complicating practical implementation of large-scale scp-MS investigations. Data-dependent acquisition (DDA) based methods have dominated the field thus far, led by the development of SCoPE-MS approach^4,10,11,14^. The method utilizes isobaric TMT labelling to multiplex single-cells and combines them with a carrier channel containing 100-200 cells, allowing parallel analysis of up to 16 cells in a single run with the latest TMTPro 18-plex reagent set. This tremendously improved the throughput and proteome coverage of scp-MS, but in-depth explorations of the biases introduced by the carrier channel in terms of protein quantification have clarified the benefits and limitations of this method^7,10,15,16^. Latest label-free quantification (LFQ) - based approaches have significantly improved the proteome coverage (1000-2000 proteins) and surpass DDA multiplexing based workflows, although the low throughput remains a significant challenge^12,17^. A dual-column LC configuration has been proposed as a potential solution, but is yet to be demonstrated on actual single-cell input^18^. Data-independent acquisition (DIA) ^19,20^ based approaches have also been used to tackle single-cell proteomes and currently provide the deepest proteome coverage^3,6,21^. Furthermore, the introduction of plexDIA increased the throughput by allowing single-cell multiplexing, similarly to SCoPE-MS, demonstrating great potential for increased throughput in DIA-based approaches^6^.

Due to the ultra-low amount of peptides derived from a single-cell, long injection times (ITs) are required to ensure sufficient ions are collected for identification and quantification^7,11,12,15,22^. This limits the capacity of DDA based methods to comprehensively sequence all the peptides present in the sample, putting great demands on analysis efficiency in terms of effectively using available mass analyzer time (Furtwängler et al., 2022; Huffman et al., 2022). In contrast, DIA does not suffer from such limitations as multiple peptides are co-isolated and analyzed, potentially acquiring both the MS1 and MS2 spectra of all the precursor ions present in the samples^23^. However, identification and quantification can be hindered by spectra convolution and low signal intensity. Improvements in chromatographic separation have the potential to benefit all types of scp-MS workflows, by providing higher resolution (sharper peaks boosting peptide ion flux), better separation capacity and more stable retention times run-to-run. Accordingly, narrow-bore columns and perfectly ordered micropillar array-based nano-HPLC cartridges (uPAC) have been manufactured and have shown promising results for ultra-low (<1ng) input proteomics^17,24–26^. uPAC columns have shown great promise for low-input (<10ng) proteomics, with high separation power and exceptionally robust peptide retention times^24,25^. Impressively, the improvements brought about by the uPAC columns allowed quantification of proteins from only 50pg of input^26^.

DIA holds great promise for scp-MS and low-input proteomics, however optimal method designs with regards to input load have not been comprehensively investigated. In this study, we carry out survey experiments to determine to which extent optimal DIA method designs are dependent on the sample input load. We build further on our findings by utilizing a high-resolution MS1 (HRMS1)-based DIA approach, to generate a new low-input DIA method design, which we combine with the newly developed uPAC Neo limited sample analytical column. We showcase that with a combination of advanced data acquisition and latest-generation chromatography, we can obtain proteome coverage from low-input (10ng) samples that is reminiscent of standard (100ng) samples. A strong focus throughout this work was on keeping sample throughput high, and therefore we opted to assess short gradients only, as implemented either on an Ultimate3000 with flowrate-ramping, or an EvoSep One chromatography system for the initial DIA scheme evaluations. We epitomize our combined findings, by profiling single-cell proteomes with the use of gas-phase fractionated (GPF) libraries generated from single-cells, to establish a new workflow that can be used for deep proteome quantification with a high degree of data completeness.

## 2. Methods

### Cell culture and FACS sorting

HEK cells were cultured in RPMI media containing 10 % FBS and 1 % Penstrep. Upon reaching 80 % confluence, cells were harvested and washed with ice-cold PBS to remove any remaining growth media prior FACS sorting and finally resuspended in ice-cold PBS at 1e6 cells/ml. Cell sorting was done on a FACS Aria III instrument, controlled by the DIVA software package (v.8.0.2) and operated with a 100 *μ*m nozzle. Cells were sorted at single-cell resolution, into a 384-well Eppendorf LoBind PCR plate (Eppendorf AG) containing 1 *μ*L of lysis buffer (100 mM Triethylammonium bicarbonate (TEAB) pH 8.5, 20 % (v/v) 2,2,2-Trifluoroethanol (TFE)). Directly after sorting, plates were briefly spun, snap-frozen on dry ice for 5 min and then heated at 95 °C in a PCR machine (Applied Biosystems Veriti 384-well) for an additional 5 mins. Samples were then either subjected to further sample preparation or stored at −80 °C until further processing.

### Sample preparation of single cells for mass spectrometry

Single-cell protein lysates were digested with 2 ng of Trypsin (Sigma cat. nr. T6567) supplied in 1 *μ*L of digestion buffer (100 mM TEAB pH 8.5, 1:5000 (v/v) benzonase (Sigma cat. nr. E1014)). The digestion was carried out overnight at 37 °C, and subsequently acidified by the addition of 1 *μ*L 1 % (v/v) trifluoroacetic acid (TFA). The resulting peptides were either directly submitted to mass spectrometry analysis or stored at −80 °C until further processing. All liquid dispensing was done using an I-DOT One instrument (Dispendix).

### Liquid chromatography configuration

The Evosep one liquid chromatography system was used for DIA isolation window survey (Figure 1) and HRMS1-DIA (Figure 2) experiments. The 31 min or 58 min gradient, where peptide elution is carried out with 100nl/min flow rate. A 15cm × 75 *μ*m ID column (PepSep) with 1.9 *μ*m C18 beads (Dr. Maisch, Germany) and a 10 *μ*m ID silica electrospray emitter (PepSep) was used. Mobile phases A and B were 0.1% formic acid in water and 0.1% in Acetonitrile. The uPAC Neo limited samples column connected to the Ultimate 3000 RSLCnano system via built-in NanoViper fittings, and electrically grounded to the RSLCnano back-panel. For the single-column scheme the column was connected according to the “Ultimate 3000 RSLCnano Standard Application Guide” (page 38) and the autosampler injection valve, configured to perform direct injection of 1 *μ*L volume sample plugs (1 *μ*L sample loop–full loop injection mode). The pre-column scheme was also assembled according to the Standard Application Guide (page 47). The analytical column was kept in a column oven and kept a constant temperature of 40 °C. All the used gradients are available in the supplemental information. Both LC systems were coupled online to an orbitrap Eclipse Tribrid Mass Spectrometer (ThermoFisher Scientific) via an EasySpray ion source connected to a FAIMSPro device.

**Figure 1.**
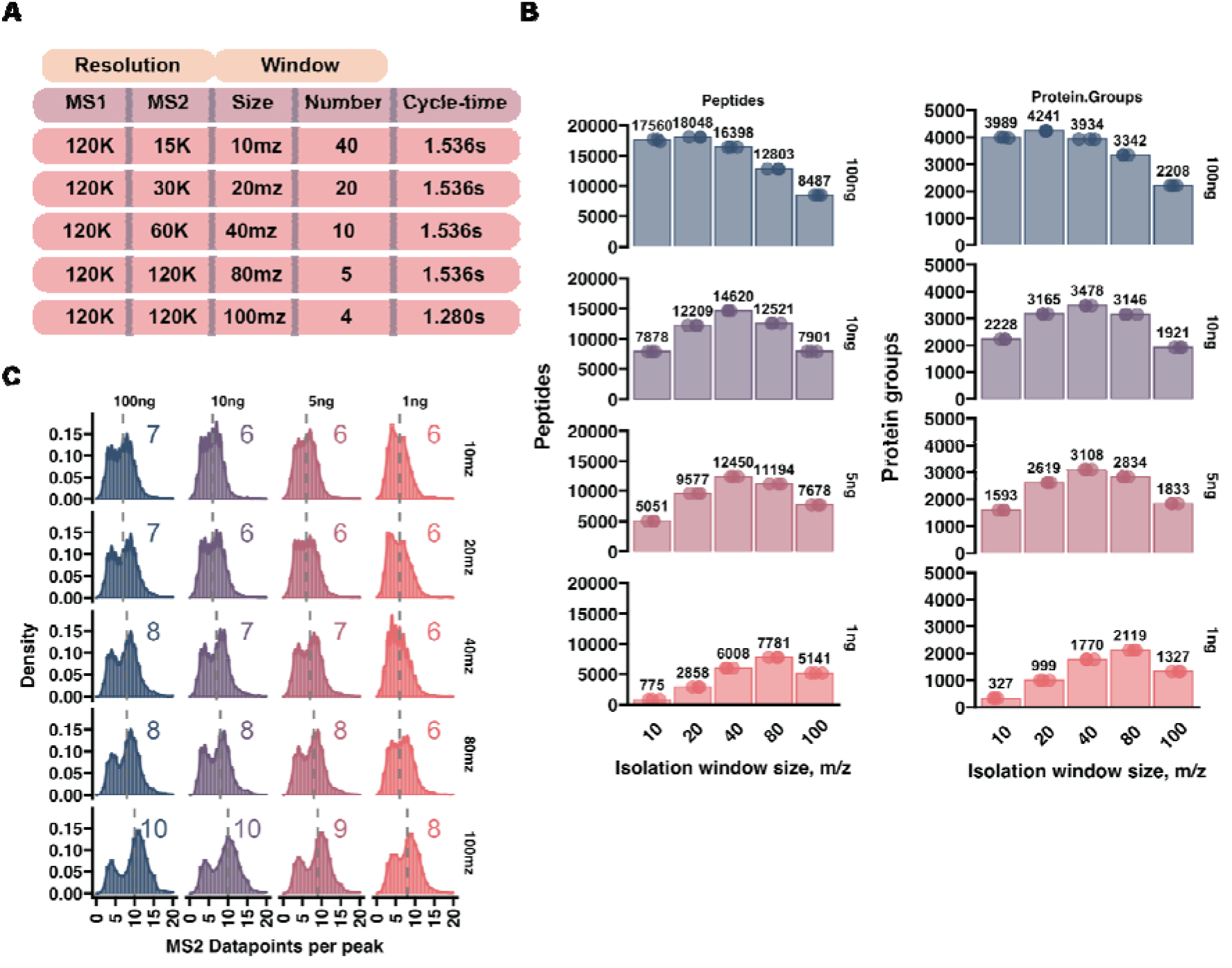
Optimal DIA window isolation size is dependent on the amount of input material. **A)** Table summarizing the different DIA acquisition methods used. **B)** Barplot of peptide and protein quantification numbers with DIA methods that have varying isolation window size and resolution. **C)** Histograms of points-per-peak quantified for peptides. Only one replicate out of three is shown. Grey dashed line marks the median PPP, the number of which is listed at the top of each to the histogram.

**Figure 2.**
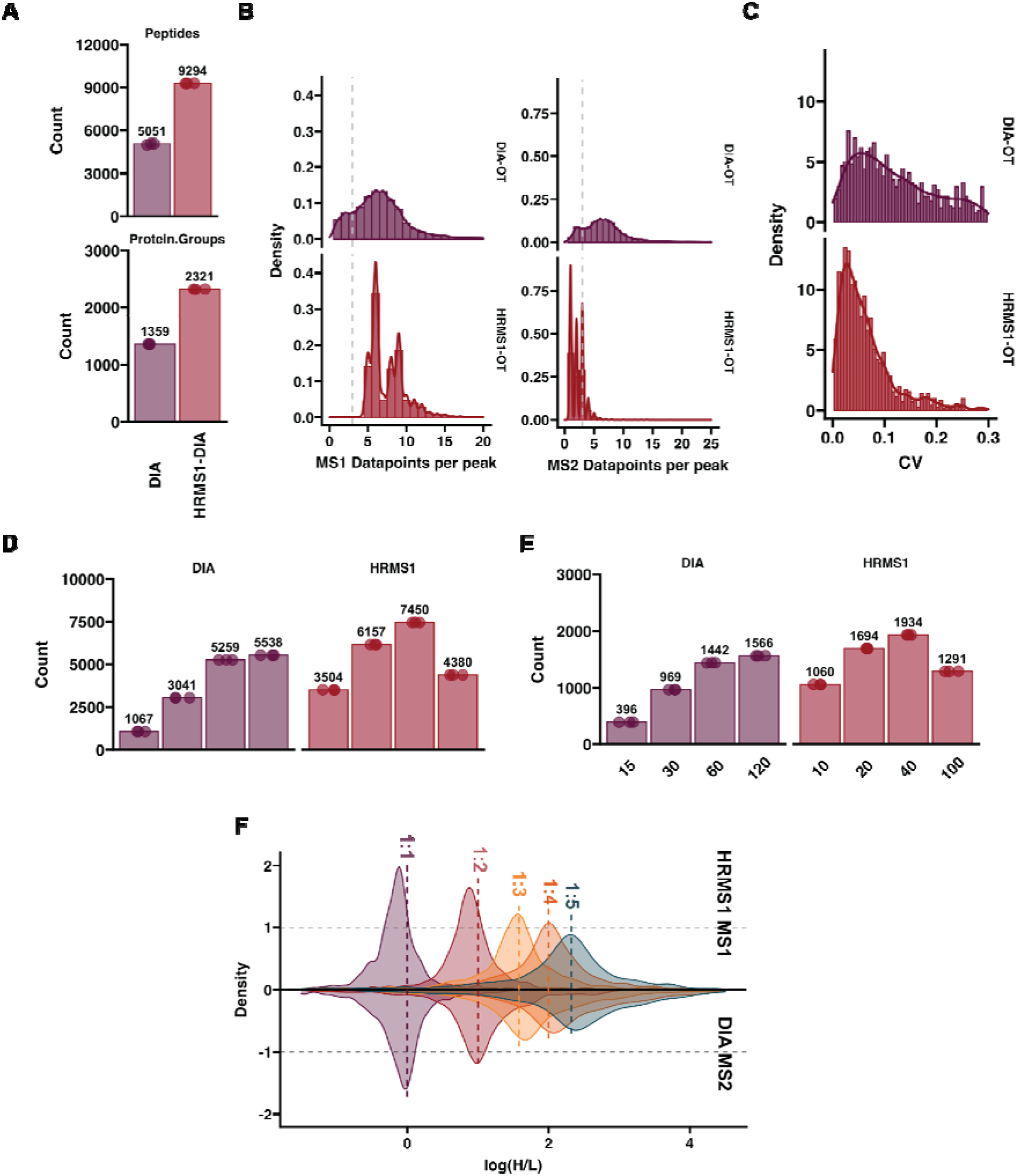
WISH-DIA enables deeper proteome profiling from low-input. **A)** Quantified number of peptides and proteins with standard DIA and HRMS1-DIA with 1ng injection with 20PSD Evosep whisper method. **B)** Acquired MS1 and MS2 points-per-peak. **C)** CV distribution histograms of quantitative precision of standard and HRMS1 DIA. Respectively, MS2 and MS1 based quantification was used for CV comparison. **D-E)** Barplots of quantified peptides and proteins in wide window HRMS1-DIA acquisition survey with 40SPD Evosep whisper method from. **F)** Heavy and light protein abundance ratio density plots. 1ng total input material was kept constant and the amount of light and heavy peptides were varied to achieve the required ratios. HRMS1 MS1 based quantification is shown in the top, and DIA MS2 based quantification in the negative side of the plot.

### MS data acquisition

The mass spectrometer was operated in positive mode with the FAIMS Pro interface compensation voltage set to −45 V. Different DIA acquisition methods were used and are outlined in the results section. Albeit certain constant parameters were used. The MS1 scans were carried out at 120 000 resolution with an automatic gain control (AGC) of 300% and maximum injection time set to auto. For the DIA isolation window survey a scan range of 500-900 was used and 400-1000 rest of the experiments. Higher energy collisional dissociation (HCD) was used for precursor fragmentation with a normalized collision energy (NCE) of 33% and MS2 scan AGC target was set to 1000%.

### Data analysis

Spectronaut 16 and 17 (pre-release) versions were used to process raw datafiles. Direct DIA analysis was run on pipeline mode using modified BGS factory settings. Specifically, the imputation strategy was set to “None” and Quantity MS level was changed to MS1. Trypsin and LysC were selected as digestion enzymes and N-terminal protein acetylation and methionine oxidation were set as variable modifications. Carbamidomethylation of cysteines was set as fixed modification for experiments that used diluted Hela peptides and removed when single-cell runs were searched. The single-cell GPF library runs were added to directDIA to supplement the single-cell dataset search. SILAC experiments were processed in Spectronaut 16, with the Pulsar search engine setting altered to accommodate multiplexed samples. Two label channels were enabled and fixed Arg10 and Lys8 modifications were added to the second channel. The in-Silico Generate Missing channel setting was used with the workflow set to “label. The complete Spectronaut settings can be downloaded from the MassIVE repository (see Data availability). Protein and peptide quantification tables were then exported and analyzed in R (version 4.2.2) in the Visual Studio Code editor environment (version 1.73), with additional packages: tidyverse (doi: 10.21105/joss.01686), and ggprism (doi: 10.5281/zenodo.4556067).

### Hela tryptic digest preparation

Cells were harvested at 80% confluence and lysed in 5% sodium dodecyl sulfate (SDS), 50mM Tris (pH 8), 75mM NaCl, and protease inhibitors (Roche, Basel, Switzerland, Complete-mini EDTA-free). The cell lysate was sonicated for 2 × 30s and then was incubated for 10 min on ice. Proteins were reduced and alkylated with 5mM tris(2-carboxyethyl)phosphine (TCEP) and 10mM CAA for 20 min at 45 °C. Proteins were diluted to1%SDS and digested with MS grade trypsin protease and Lys-C protease (Pierce, Thermo Fisher Scientific) overnight at an estimated 1:100 enzyme to substrate ratio quenching with 1% trifluoroacetic acid (TFA) in isopropyl alcohol. For the cleanup step by styrenedivinylbenzene reverse-phase sulfonate (SDB-RPS)^27^, 10 *μ*g of peptides was loaded on StageTip^28^ and washed twice by adding 100 *μ*L of 1% TFA in isopropyl alcohol. Peptides were eluted by adding 50 *μ*L of an elution buffer (1% Ammonia, 19% ddH2O,and 80% Acetonitrile) in a polymerase chain reaction (PCR) tube and dried at 45 °C in a SpeedVac. Lastly, peptides were resuspended in buffer A and their concentration was measured by nanodrop.

### Data availability

The complete MS raw data, Spectronaut search files have been deposited to MassIVE under the following accession MSV000090792.

### Code availability

The code used to generate to process the tables exported from Spectronaut analysis has been stored in the following repository: https://github.com/NotValdemaras/uPAC-Neo

## 3. Results

### Increasing low-input sample proteome coverage by wide DIA isolation windows

Increasing the isolation window size during DIA based acquisition should in theory hamper peptide identification due to more extensive precursor co-isolation resulting in increasingly chimeric spectra. While this effect is pronounced for high-load (>10ng) samples, we hypothesized that co-isolation constraints are not as prevalent when handling low-load samples (<10ng). To test this, we carried out a series of experiments where we injected different amounts of Hela digest (100ng, 10ng, 5ng and 1ng) and acquired the MS spectra with DIA methods of varying isolation window sizes and resolutions combined with varying ion ITs, while maintaining approximately the same scan-cycle time (Figure 1A). As expected, 100ng of input material resulted in the highest number of protein identifications. Doubling the isolation window width from 10m/z to 20m/z, and doubling the resolution slightly increased the proteome coverage, however further widening beyond 20m/z had an opposite effect (Figure 1B). In contrast, when lower amounts of peptide were injected, 40m/z isolation window gave the best results for 10 and 5 ng. Decreasing the peptide load to 1ng further moved this optimal value to 80m/z (Figure 1B), suggesting that the chimeric spectra effects due to co-isolation at such loads are sufficiently low to be overcome by increased resolution and IT. All the methods had a median of 6 or more points-per-peak to ensure comparable quantification potential (Figure 1C). Interestingly, although the scan cycle-time was kept constant, increasing the resolution, isolation window size, and ITs, led to more data points-per-peak (Figure 1B). Accordingly, protein quantification precision also improved as more data points were collected, which was especially marked at the lowest-level 1ng injections (Figure S1A). The additional points are detected potentially due to longer ITs which allows quantification of the elution profile tails that fall below the background intensity at shorter ITs. Together, these findings indicate that chimeric spectra effects are not as pronounced in low-input samples and can be overcome sufficiently by increased resolution/ITs, facilitated through wider DIA isolation windows.

### HRMS1-DIA in combination with wide isolation windows enhanced quantified proteome depth

Since DIA acquires both MS1- and MS2-level spectra, quantification can be carried out on either level, with the latter commonly being attributed to be more accurate in the literature, as it can overcome co-elution biases^29,30^. Due to this, MS2-based quantification is generally preferred in DIA experiments and is the default output by most popular search engines, such as Spectronaut and DIA-NN^29,31^. A method that breaks away from this convention has also been proposed, termed high-resolution-MS1 (HRMS1) DIA^32^. While in standard DIA, the MS1 scan is followed by MS2 scans that sequentially measure the whole m/z range of interest, HRMS1 slices the total m/z range into smaller segments, interjecting MS1 scans in between (Figure S2A). This modification drastically decreases the amount of MS2 data points acquired for each precursor, eliminating the ability to perform robust quantification on the fragment level. Quantification becomes primarily focused on the MS1 information, while the MS2 is used only for identification. Accordingly, the available cycle time can now be more optimally used for a segmented part of the overall m/z range, affording longer ITs and higher resolution (Figure S2A). We compared standard DIA versus HRMS1 to determine if we can further increase our proteome coverage with this method. By modifying the DIA acquisition method according to HRMS1, we could increase our resolution (and corresponding ITs) from 30K to 60K and decrease our isolation window size from 15m/z to 8m/z, while maintaining identical scan cycle-times. Not only did HRMS1 significantly outperform standard DIA in terms of identification (Figure 2A), it also collected more points-per-peak (Figure 2B) which translated into higher quantitative precision (Figure 2C). The extra identifications by HRMS1 primarily arose from low-abundant proteins (Figure S2B). We also adopted this modification to linear ion trap (LIT) based DIA^33,34^ and observed similar overall performance gains (Figure S3A-C), although it did not surpass OT-based HRMS1-DIA.

We performed a similar isolation window survey experiment as above to see if we could synergize the HRMS1 method with wide isolation windows. In line with our initial observations, widening the isolation window to accommodate for longer ITs and higher resolution scans on 1ng injections resulted in increased numbers of quantified proteins (Figure 2D). The protein count peaked at 40m/z isolation width and decreased once 100m/z was reached. We term our tailored low-input method WISH-DIA (**W**ide **IS**olation window **H**igh-resolution MS1-DIA), to epitomize the combination of wide isolation windows and use of HRMS1 quantification.

Although WISH-DIA showed great promise, the question of quantitative bias remained due to MS1-based quantification. To evaluate this aspect, we utilized a SILAC approach and mixed peptides derived from Hela cells cultured in light or heavy media in different ratios and analyzed the data with the best performing methods (Figure 2E). While keeping the total sample load to only 1ng to carefully mimic a low sample-load setting, we directly compared protein abundance (H/L) ratios derived from DIA fragment level or HRMS1 precursor-level (Figure 2F). Both showed a ratio distribution that was in line with the expected values. There was a clear drop in accuracy as the ratio of light and heavy peptides was increasing, potentially, due to the decreasing proportion of light peptides in the samples making them harder to quantify. MS1 yielded sharper peaks compared to MS2, indicating higher quantitative accuracy, albeit a minor, but clear bias could be observed when 1:1 and 1:2 mixtures were compared on MS1 level quantification, which was not present when MS2 was used. Interestingly, when higher ratio mixtures were compared, there appeared a minor, but clear discrepancy in MS2 level quantification, while MS1 ratio distribution remained centered around the expected value (Figure 2F). Higher MS1 accuracy was also observed comparing MS1 and MS2 protein ratios from the standard DIA method (Figure S3D). Taken together, we conclude that WISH-DIA enhances proteome depth from low-input samples while maintaining robust quantitative accuracy.

### Micropillar array-based nano-HPLC cartridges/columns for low-input proteomics

Next, we substituted the packed C18-beads column with a next-generation *μ*PAC Neo Low Load column to further augment our low-input workflow efforts (Figure 3A). This 50cm column has a reduced cylindrical pillar diameter of 2.5μm, an interpillar distance of 1.25μm, a total column volume of 1.5μL, and is non-porous, thereby increasing its chromatographic performance at much reduced loading capacities. We designed methods that utilized flow-ramping up to 500 nl/min to minimize the overhead time needed for peptide break-through and analytical column regeneration (Figure 3B). We generated three single-column and two pre-column configuration methods and tested chromatographic performance of the column by running tryptic digests with our developed WISH-DIA methods (Figure 3C). Examining the peak-width of the single-column configuration, we saw that the full-width at half maximum (FWHM) of the peptide precursors peaks is approximately 6.6 second, which broadened to 8.58 seconds for the longest method in line with total gradient time (Figure 3D). Addition of a pre-column in-line resulted in increased peak-widths > 9 seconds, however extending the gradient only resulted in a marginal increase in peak-width (Figure 3D). Retention times were very robust across runs, with almost all precursor elution apex deviations being limited to 2.5 seconds (Figure 3E), underlining the solid chromatographic performance of this novel *μ*PAC Neo Low Load column. We proceeded to further benchmark the analytical column in terms of proteome coverage for variable amounts of input material.

**Figure 3.**
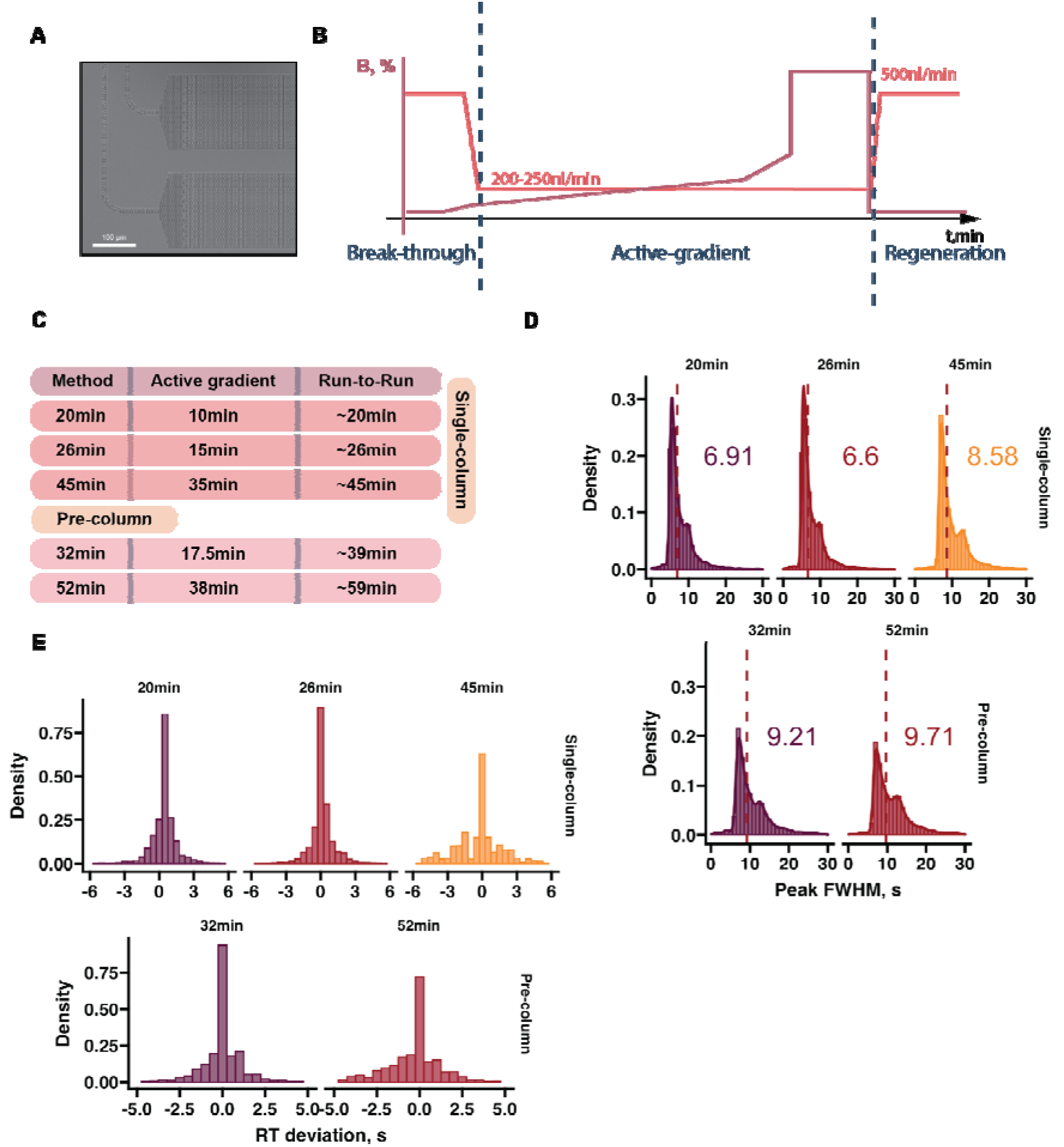
Chromatographic performance of the uPAC Neo limited-sample analytical column. **A)** Scanning electron microscopy (SEM) image of the micropillar-column. **B)** Schematic visualizing the gradient used with the uPAC column, including flow-ramping capability of the Ultimate3000. **C)** Table summarizing the used methods. **D)** Histograms of peak FWHM with different method lengths with the single-column or pre-column configuration. **E)** Histogram showing the peptide elution peak apex deviation from a mean calculated from three replicates. The deviations of one replicate from the mean are shown. The plotted peak parameters were obtained from 5ng DIA runs analyzed with Spectronaut16.

### Utilizing the synergy between *μ*PAC Neo Low Load and wide isolation window HRMS1-DIA for low-input proteomics

To date, the vast majority of low or ultra-low level input (≤250pg) studies have focused on DDA based acquisition. It is now possible to routinely quantify > 1000 protein groups from such amounts ^12,25,35–37^. However, this tends to require long LC-MS instrument run-times (>1h), unless a double-barrel approach is used^18^. First, to try and maximize sample throughput, we evaluated the performance of 45, 26 and 20 minute methods (32, 55 and 72 samples per day (SPD) respectively) and injected different amounts of digested peptide in a single-column configuration (Figure 3C). Commercially available Pierce Hela digest was used (Part #88328), to ensure that our reported performance numbers can be easily evaluated by others. To fully realize the potential of the *μ*PAC Neo Low Load column, we utilized WISH-DIA to quantify proteomes from low-input material (<10ng). From 10ng we quantified from 3000 to 4700 protein groups depending on the method used (Figure 4A). Decreasing the amount of input material resulted in fewer protein identifications, albeit up to ~ 4000 and ~ 3000 protein groups could still be quantified from 5 and 1 ng. At ultra-low input level of 250pg, we quantified 2089 protein groups on average at 32SPD and 1461 at 72SPD. Overall, our workflow quantifies PG numbers comparable to previously published work, however at 2-3 times greater throughput^17,18,26,38^.

**Figure 4.**
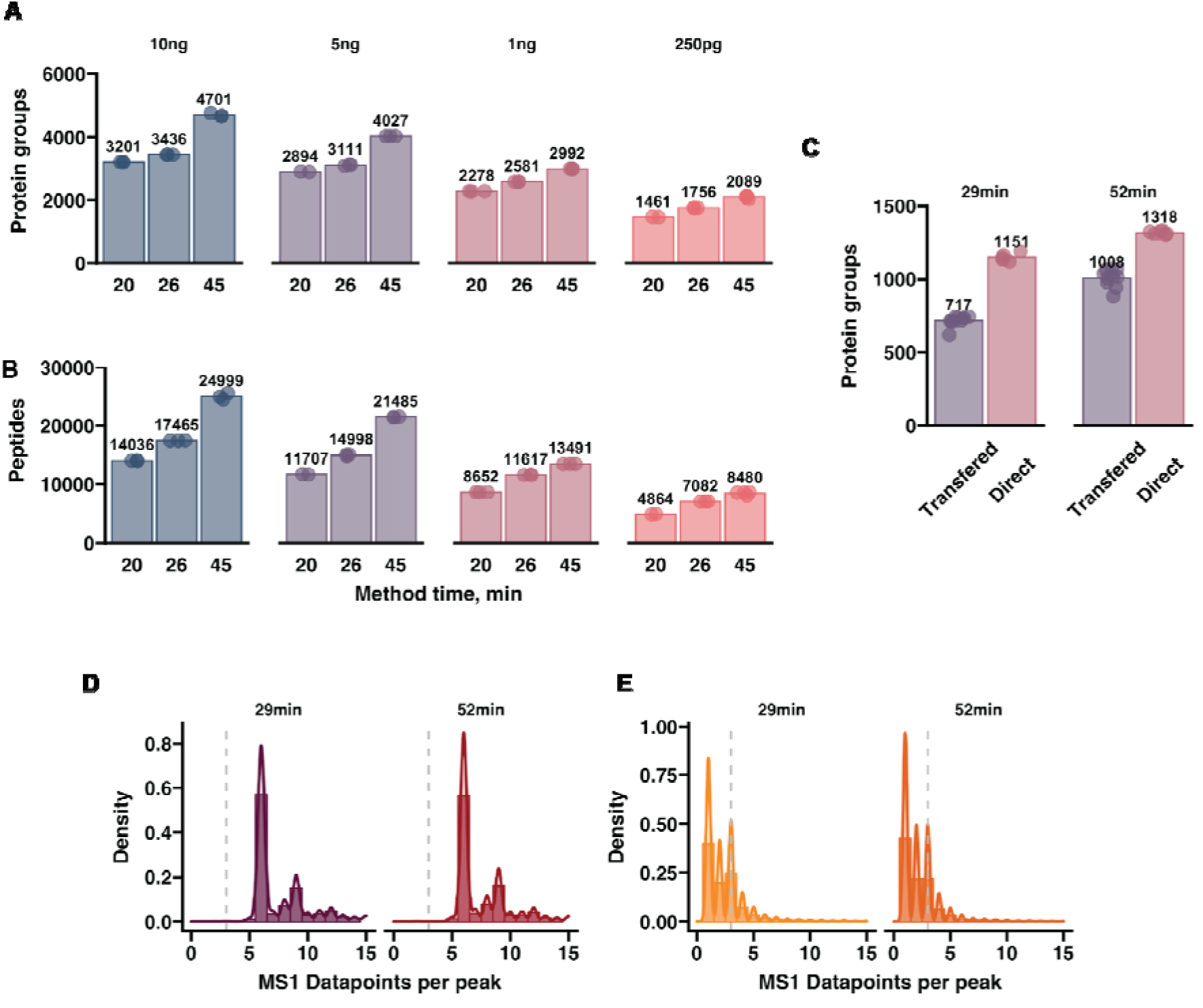
Deep proteome coverage of low-input samples by advances in multiple aspects of mass spectrometry. **A**) Barplots of quantified peptides and proteins width different method lengths and peptide loads with the single-column PAC Neo Low Load set-up. **B**) The same as A, but with the uPAC pre-column in-line. **C**) Shorter methods with the uPAC pre-column setup with 1ng of input material. **D**) The same as C, however, column regeneration was parallelized with pre-column loading, shortening the 32 and 23 min gradients to 17 and 27 min, respectively. **E**) Barplot showing quantified peptides and proteins from single-cell input (1 HEK293 cell). Two methods 120k-18w-34mz and 240k-9w-68mz. 52min gradient was used. Cells were transferred to a 96 well plate before injection. **F**) Cells directly injected from a 384 well plate with two gradient lengths, 240k-9w-68mz method. All reported numbers are obtained with directDIA by searching the runs from the same method in a single batch. Peptide identification numbers are provided in Figure S4.

To process biologically relevant samples where standard solid-phase extraction ^28^ cannot be used, a pre-column can be used to ensure robustness on the chromatographic system, and prevent clogging by non-protein contaminants present in the samples. This is especially relevant in single-cell proteomics^1^ where indeed prior sample clean-up is not possible. With a tailor-made *μ*PAC pre-column setup, consisting of non-porous 5*μ*m pillars based on C8, we developed 32 and 52 minute methods that could quantify similar peptide and protein group numbers as a single-column setup (Figure S4A-B). Due to the larger sample loop used (20ul vs. 1ul in the single-column setup), the pre-column configuration adds 7 min overhead time to each method, decreasing throughput to 36 and 24 SPD (Figure 3C). With the pre-column configuration, we achieved reminiscent proteome coverage compared to the single-column set-up, where we could quantify > 2000 protein groups from ultra-low input (Figure S4A-B). This was slightly unexpected as the pre-column leads to peak broadening (Figure 3D). The deep coverage benefited not only from improved acquisition and chromatography methods, but also the computational advance of Spectronaut17 (Bruderer et al., 2015), which increased the identification of ultra-low input by ~ 30%, relative to the previous version (Figure S4A-B). As the ultimate goal of our work was to be able to analyze single-cell proteomes, both with high proteome depth and quantitative accuracy, and at reasonable throughput, we next evaluated the performance of WISH-DIA on actual single cells. HEK293 cells were prepared in 384 well Eppendorf low-bind plates with previously described protocols (See methods) and transferred to a 96 well plate for injection. Since single-cell samples have been shown to require high ITs ^7,11,12,15,22^, to accommodate this we further increased the IT and resolution of our WISH-DIA method from 120k (246ms IT) to 240k (502ms IT), while doubling the isolation window size (68 m/z) to maintain the same scan cycle time (Figure S4C). We processed 10 single-cells with our two established 29min and 52min precolumn methods and could quantify 717 and 1008 proteins by directDIA (Figure 4C). However, as also recently shown by others^35^, transferring single-cell samples leads to severe signal losses. To test the extent of this effect in our experimental setup, we directly injected the peptides derived from single cells from their original 384-well plate. Accordingly, direct injection boosted our average identifications by ~ 60% for the shorter and ~ 30% for the longer method (Figure 4C), bringing our quantified protein numbers to 1151 and 1318 when searched with directDIA. Quantification robustness was ensured by keeping the cycle time sufficiently short to collect a minimum of 5 data points per precursor elution profile (Figure 4D), while MS2 data points were only collected for identification (Figure 4E).

### Quantification quality of additional proteins gained by high-load library use

Some studies have chosen to utilize enhanced search strategies by including higher load libraries (e.g. 10ng), which can drastically boost the number of quantified proteins. So far, either diluted bulk cell population digests or samples containing multiple cells have been used for this purpose^3,35,39,40^. However, the exact impact of using such high-load (HL) ID transfer approaches remains unclear, especially in terms of quantification accuracy, as peptides that are potentially lost during single-cell processing might be erroneously quantified. A gas-phase-fractionated library (GPF^41,42^) is another approach that can be used to gain identifications, which is generated by dividing our m/z range of interest into 6 segments of 100m/z and analyzing samples while acquiring spectra for only that segment (See Methods). Due to the decreased m/z range for each individual run, we could therefore further increase our ITs (1014ms) and decrease the isolation window width, allowing the identification of peptides that have very low abundance and are difficult to quantify in our global WISH-DIA runs.

To assess the protein quantification quality of both approaches, we mixed light and heavy peptides in three different ratios while maintaining a constant 10ng injection load. We then diluted our sample to 1ng injections that were used as actual runs and GPF library creation, and the 10ng were used to acquire HL libraries. To gauge the quantification accuracy we plotted the light and heavy ratio distributions for the identified proteins obtained with directDIA or LibraryDIA with a high-load or GPF library (Figure 5A). The use of a HL library approximately doubled the coverage, while GPF led to ~ 50% additionally identified proteins. The enhanced proteome depth was accompanied by substantial widening of the ratio distribution, indicating loss of accuracy in the dataset as a whole (Figure 5A). This was primarily driven by the addition of a large number proteins with low quantitative accuracy (Figure 5B). The quantitative accuracy of the proteins that could be identified by directDIA remained similar, however some detrimental effects due to the library search can be noted, likely due to the detection of additional low-abundant peptides (Figure S5).

**Figure 5.**
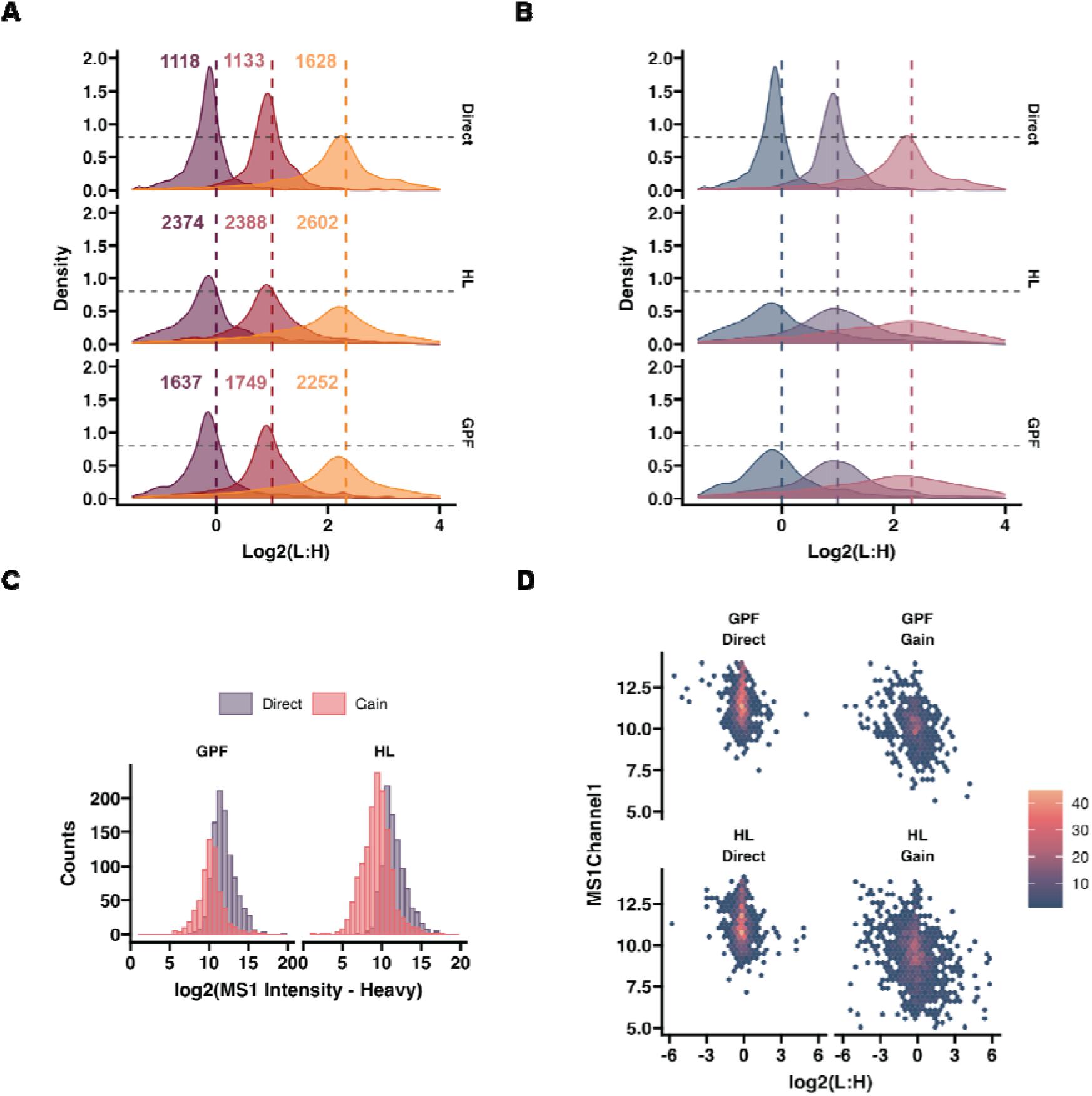
Assessing the quality of protein quantification gained by high-load and gas phase fractionated library based DIA. **A)** Density plots showing the log2 transformed light and heavy protein abundance ratios. Proteins quantified with directDIA shown in top and high-load (HL) library in the middle and gas-phase fractionated (GPF) in the bottom. Dashed lines denote expected ratios and numbers indicate the total number of identified proteins in each mix. MS1-based quantification is used throughout. **B)** Similar to A, but showing the distribution of proteins identified with directDIA and gained with either HL or GPF libraries. **C)** Histogram showing protein distribution across the log2 transformed abundance range. Only the 1:1 (L:H) mix data is shown. **D)** Hexbin plot showing the distribution of light and heavy (L:H) log2 transformed protein ratios for proteins found with directDIA and gained by the HL and GPF libraries.

As low abundant proteins are expected to naturally have poorer quantification relative to high abundant ones, we investigated this in greater detail. Application of libraries tremendously improved the identification of proteins in the lowest end of the abundance range (Figure 5C), but the gained proteins did extend beyond this range. The HL clearly aided the identification of a larger number of proteins found in the lowest end of the abundance range compared to GPF, indicating its higher capacity to extend proteome coverage. Interestingly, the light and heavy peptide ratios were more dispersed throughout the abundance range for both libraries, suggesting that the quantification quality of those gained proteins is potentially rather poor (Figure 5D). These findings point towards possible challenges with the accuracy of proteins gained via libraries from low-input samples and indicate that extra scrutiny is warranted when biologically interpreting these additional identifications.

### WISH-DIA with a next generation analytical column enables high-quality single-cell proteome profiling

As a proof of concept, we generated a small dataset of 100 cells using WISH-DIA in combination with the *μ*PAC Neo Low Load column. We analyzed 62 cells with a 40SPD method and 40 cells with 24SPD. On average, both methods quantified ~ 1670 protein groups per cell (Figure 6A). Although protein quantification was almost identical, the longer method could detect more peptides (Figure 6A). As an alternative to high-load libraries, we instead opted to generate a gas-phase-fractionated library (GPF^41,42^), by dividing our m/z range of interested into 6 segments of 100m/z and running single-cell samples while acquiring spectra for only that segment (See Methods). Due to the decreased m/z range for each individual run, we could therefore further increase our ITs (1014ms) and decrease the isolation window width, allowing the identification of peptides that have very low abundance and are difficult to quantify in our global WISH-DIA runs. By applying such a GPF approach to our single-cell runs, we were able to boost our quantified proteins by ~ 20% (Figure 6A). As expected, the quantification of these additionally identified proteins was noisier, and primarily spanned the lower range of the abundance distribution (Figure 6B). All the runs showed a relatively low level of missing values on the protein level, with the vast majority of cells exhibiting <20% missing values with directDIA (Figure 6C). However, GPF library application increased data sparsity to 30-40%. Arguably, this is an improvement for single-cell proteomics, as most studies to date have reported a high degree of missing values ~ 50%. In our case, with HEK293 being a rather homogeneous cell line, we expect that most of the variation in our data can be explained by differences in cell cycle stages. To further assess this, we integrated both the 40SPD and 24SPD datasets by standardizing the abundances and clustered the cells with both linear (PCA) and non-linear (UMAP) methods to gauge this biological variation (Figure 5D). The first principal component (PC1) captured a large degree or variation present in our dataset. To determine if PC1 was correlated with the cell cycle, we tracked the standardized abundance of the MKI67 protein, which has highest levels during G2 and mitotic cell phases. There was a clear trend as the MKI67 levels increased along the PC1 (Figure 6D). Similarly, in the UMAP analysis two clusters of cells were obtained and MKI67 levels increased along the second manifold dimension (Figure 5D). No clustering based on run order was observed, however PC2 seemed to capture method related variation, but it should be noted the percentage of variation is rather small (Figure S6A-B), underlining that our workflow can capture biologically relevant trends in single-cell proteome profiles.

**Figure 6.**
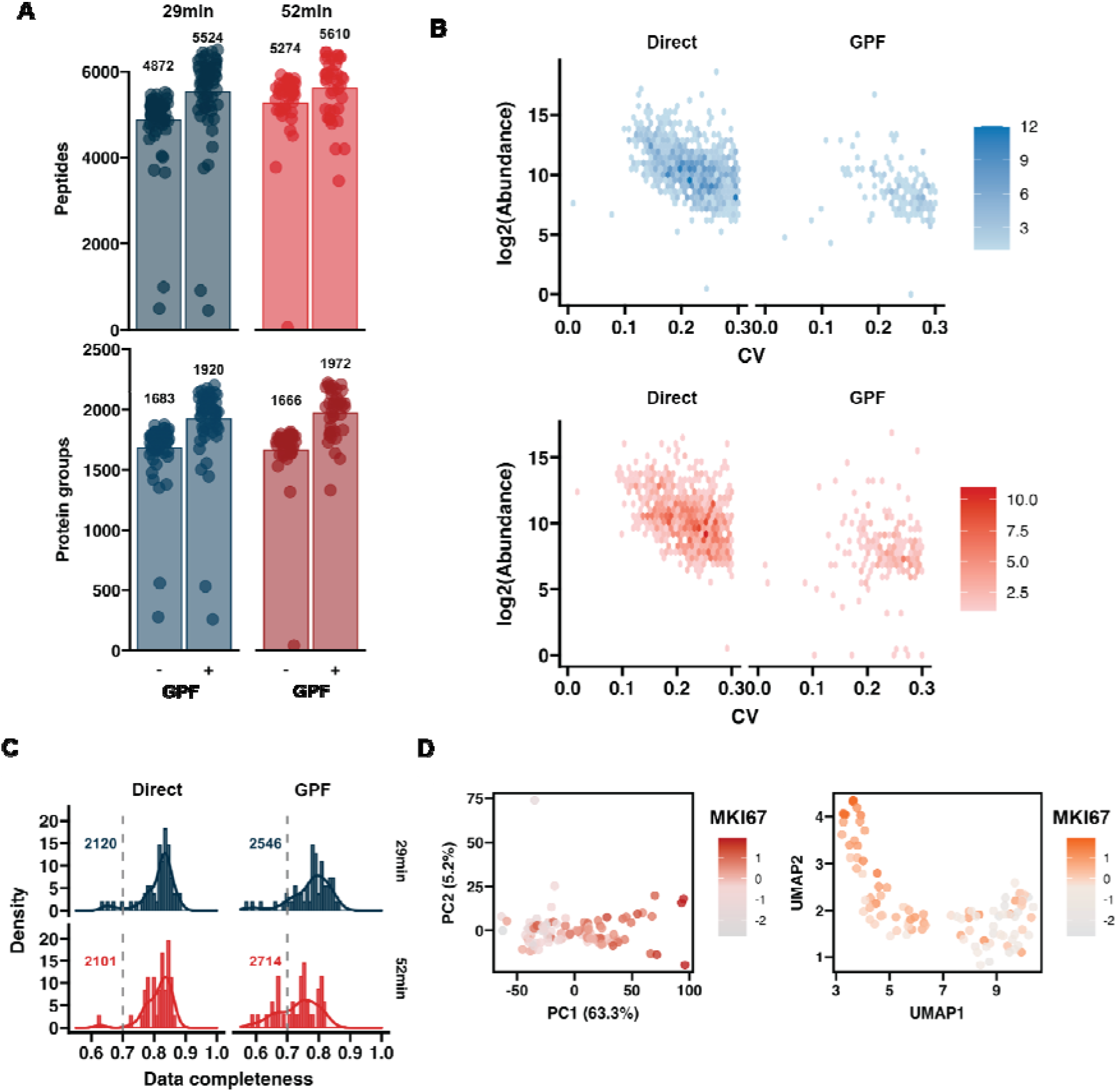
Single-cell proteome profiling. **A)** Barplots of quantified peptides and proteins from single-cell inputs with two different methods. The spectra were searched either with directDIA approach or with a GPF-based library (GPF-DIA). **B)** Hexbin plots showing the log2 transformed abundance and CV distribution for proteins quantified by directDIA and QBL. 29min in the top and 52min in the bottom **C)** Histogram of data completeness for each cell. Dashed grey lines mark 70% complete detection. **D)** Clustering of the integrated single-cell with PCA (left) and UMAP (right). Color coding denotes the standardized MKI67 protein abundance in each cell.

## Discussion

In this study, we developed a novel label-free single-cell proteomics workflow by utilizing low-input tailored DIA methods in combination with latest chromatography and computational advances. Specifically, we show that DIA method design should be adjusted accordingly to sample load for optimal performance. We discovered that for low-input samples the detrimental dynamic range and chimeric spectra effects due to large isolation windows (>20m/z) are overcome by resolution and injection time increases (Figure 1). In contrast, the same trend was not observed for high-load. We adopt a DIA approach that solely relies on precursor level quantification to further enhance sensitivity and use our findings to establish the WISH-DIA method. In tribrid instruments, the LIT can also be used to increase sensitivity while keeping isolation windows narrow^43^. We also applied the HRMS1 modification to LIT and showed that it significantly boosted proteome coverage for low-input samples (Figure S3).

By applying WISH-DIA with micropillar-array bases chromatography we were able to achieve high proteome depth for low-input samples with appropriate sample throughput. We quantified ~ 5000 protein groups from 5-10ng of input material, which is a highly relevant load for e.g. laser capture microdissection isolated tissue samples^44–46^. From ultra-low input samples (250pg) we manage to quantify > 2000 protein groups which is often considered as single-cell level input^2,3,12,18^. Such inputs generated from bulk digest dilutions are a poor proxy for true single-cell digests and numbers obtained with such samples should be interpreted with care. Accordingly, we tested our workflow with real single-cell digests and quantified ~ 2000 protein groups per single-cell at a throughput of 40 cell per day with the use of GPF libraries that boost the proteome coverage by >20% (Figure 5A). Such libraries are a great alternative to high pH for samples where offline fractionation is prohibitive, which is the case when analyzing single cells. It should be noted that the workflow uses standardized lab equipment and does not require single-cell proteomics designated liquid handling systems as in other protocols^5,8,47^, which should make the approach more accessible for general proteomics labs and facilities.

Although the throughput is lower compared to the DDA TMT multiplexing based approaches, which can analyze up to 160 cells per day at a throughput of ~ 1000 protein groups per cell^4,7,10,11^, the increased proteome depth and absences of a carrier channel and TMT quantification biases makes our LFQ workflow a solid alternative. This might be of special relevance when patient samples are of interest and collecting sufficient cells for carrier samples might not be feasible. While we ran our experiments on an Orbitrap Eclipse Tribrid instrument, it is expected that WISH-DIA methods translate directly to other Orbitrap platforms such as Exploris series instruments. Throughput can in principle be improved by adopting DIA compatible multiplexing, such as e.g. plexDIA, which has already been applied to single-cell analysis ^6^. Other DIA compatible tags, such as Ac-IP or TMT complement ion quantification could be also explored to increase throughput^48,49^. Currently, our U3000-based workflow at 40 SPD would allow one thousand cells to be analyzed within a month, which is approaching a level of maturity capable of conducting biologically relevant interrogations of heterogeneous cell systems.

## Acknowledgements

We would like to thank Robert van Ling at ThermoFisher for early access to the *μ*PAC Neo Low Load column and pre-column, and Biognosys for pre-release access to Spectronaut 17. We also thank EvoSep for elaborate collaborations on the EvoSep One instrument. Some of this work was funded by a grant from the Novo Nordisk Foundation to E.M.S. with reference number NNF21OC0071016. B.F. is the recipient of a fellowship from the Novo Nordisk Foundation as part of the Copenhagen Bioscience PhD. Programme, supported through grant NNF19SA0035442. V.P. is funded by a Leo Foundation grant awarded to S.F.T and E.M.S. (LF-OC-21-000832). P.A.F. is funded by a Danish Cancer Society grant (R324-A17978). Work in the Porse lab was supported by grants from the Svend Andersen Foundation, the Candys foundation, the Danish Cancer Society, Independent Research Fund Denmark and through a center grant from the Novo Nordisk Foundation (Novo Nordisk Foundation Center for Stem Cell Biology, DanStem; Grant Number NNF17CC0027852). U.A.D.K. acknowledges funding by a Novo Nordisk Foundation Young Investigator Award (NNF16OC0020670). We also thank all members from the Cell Diversity Lab, headed by E.M.S. for constructive input and fruitful discussions, and the DTU Proteomics Core for technical instrument support.

## Author Contributions

E.M.S. and V.P. conceived and designed the project. V.P., N.U., P.A.F., T.P and E.M.S. performed experiments, and B.F., J.O.D.B., S.F.T., U.A.D.K. and B.T.P. provided critical input. Data analysis was performed by V.P. The manuscript was drafted and revised by V.P. and E.M.S., with input from all other authors. E.M.S. supervised the work.

