## Supplementary Information for "Enhancing single-cell proteomics through tailored Data-Independent Acquisition and micropillar array-based chromatography"

### Supplemental figures

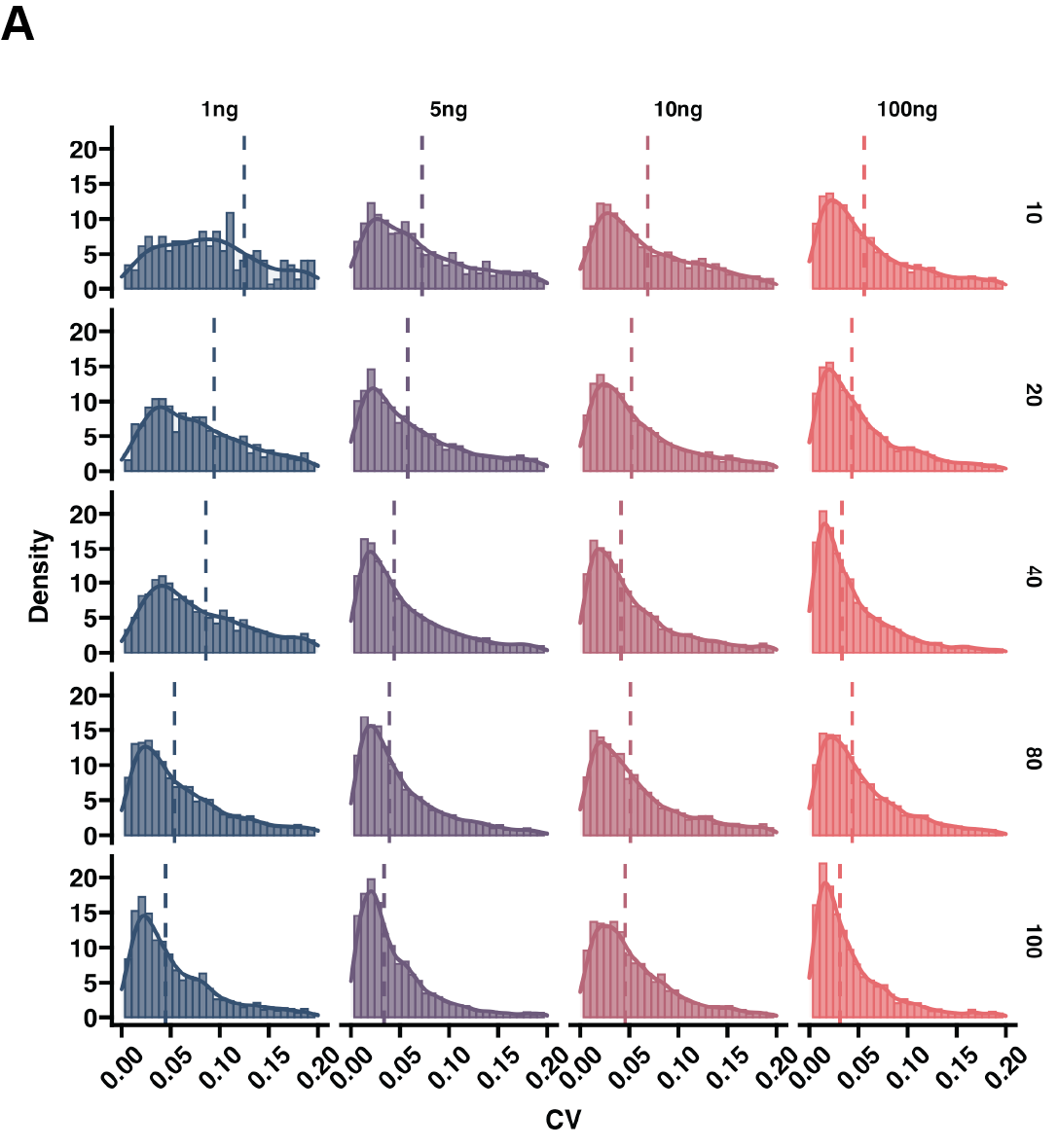

**Figure S 1.** **Higher resolution/IT improves quantification precision. A)** Coefficient of variation (CV) distribution histograms with different peptides load and DIA isolation windows. Related to Figure 1.

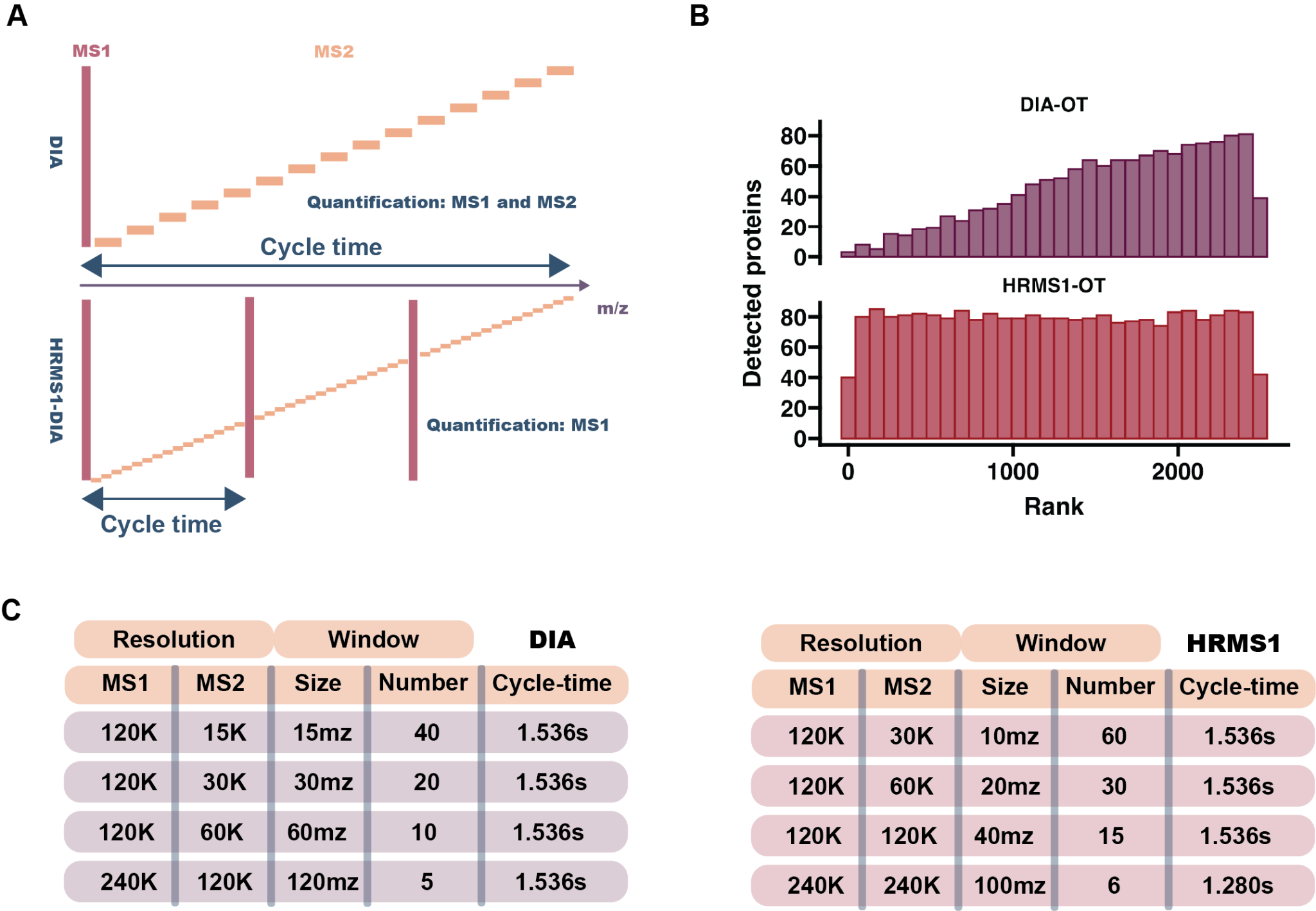

**Figure S2.** **Supporting figures and tables for HRMS1. A)** Schematic illustration of standart DIA and HRMS1 acquisition. **B)** Barplot showing detected number of proteins at a certain log transformed abundance bin with HRMS1 or standard DIA. **C)** Tables summarizing the resolution and isolation window parameters used for the window survey. Related to Figure 2.

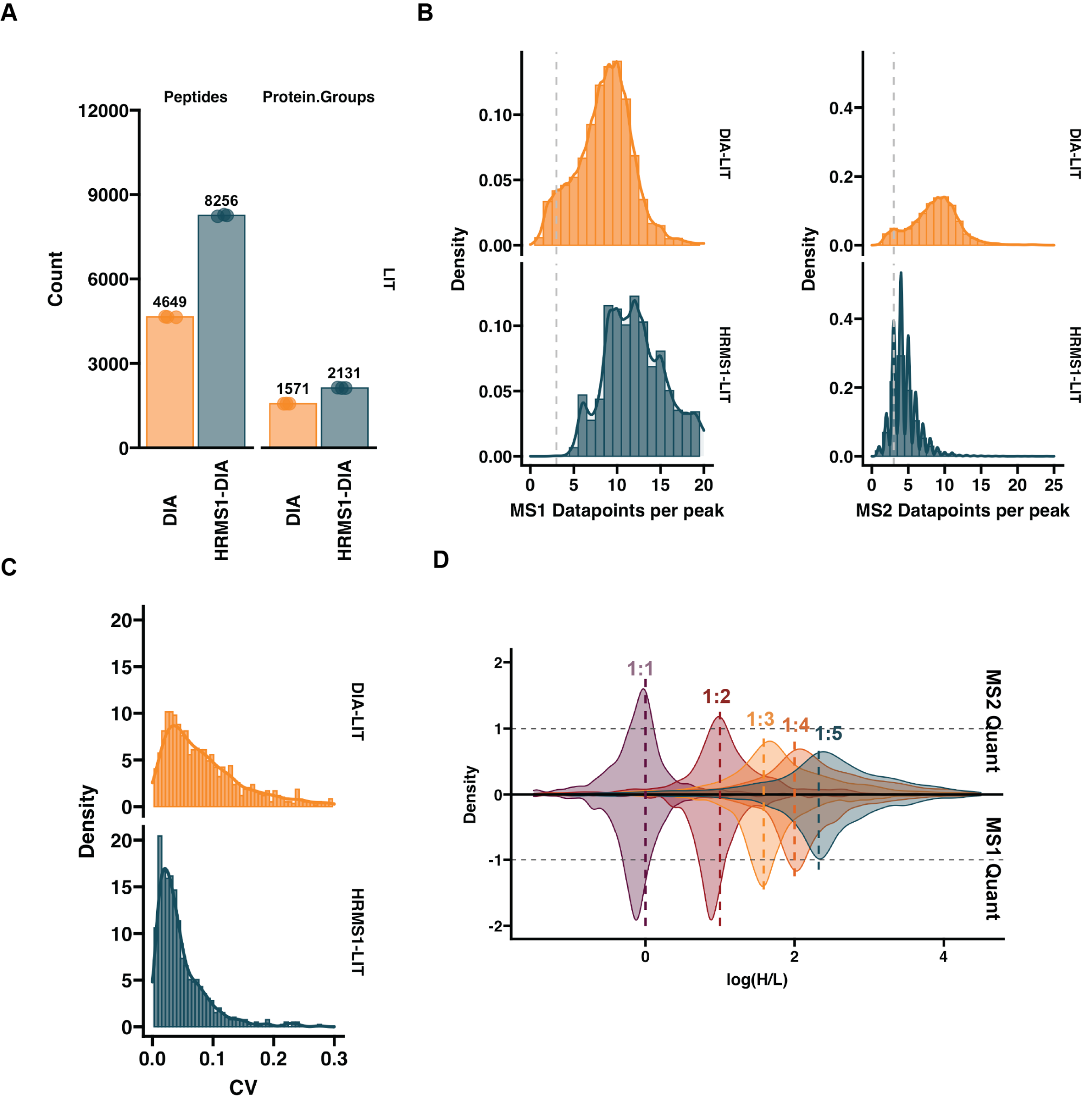

**Figure S3.** **Wide isolation window application to LIT-DIA and MS1 quantification accuracy. A)** Barplot showing detected number of proteins and peptides with LIT-DIA or HRMS1-LIT-DIA **B)** Histogram showing data points per peak on MS1 (left) and MS2 (right) level with both methods. **C)** Histogram of CV values for protein quantification. **D)** Density plots showing different SILAC light and heavy mixes. MS2 (top) and MS1 (bottom) quantification is directly compared in the plot. Dashed lines indicated expected peptide mix values. Related to Figure 2.

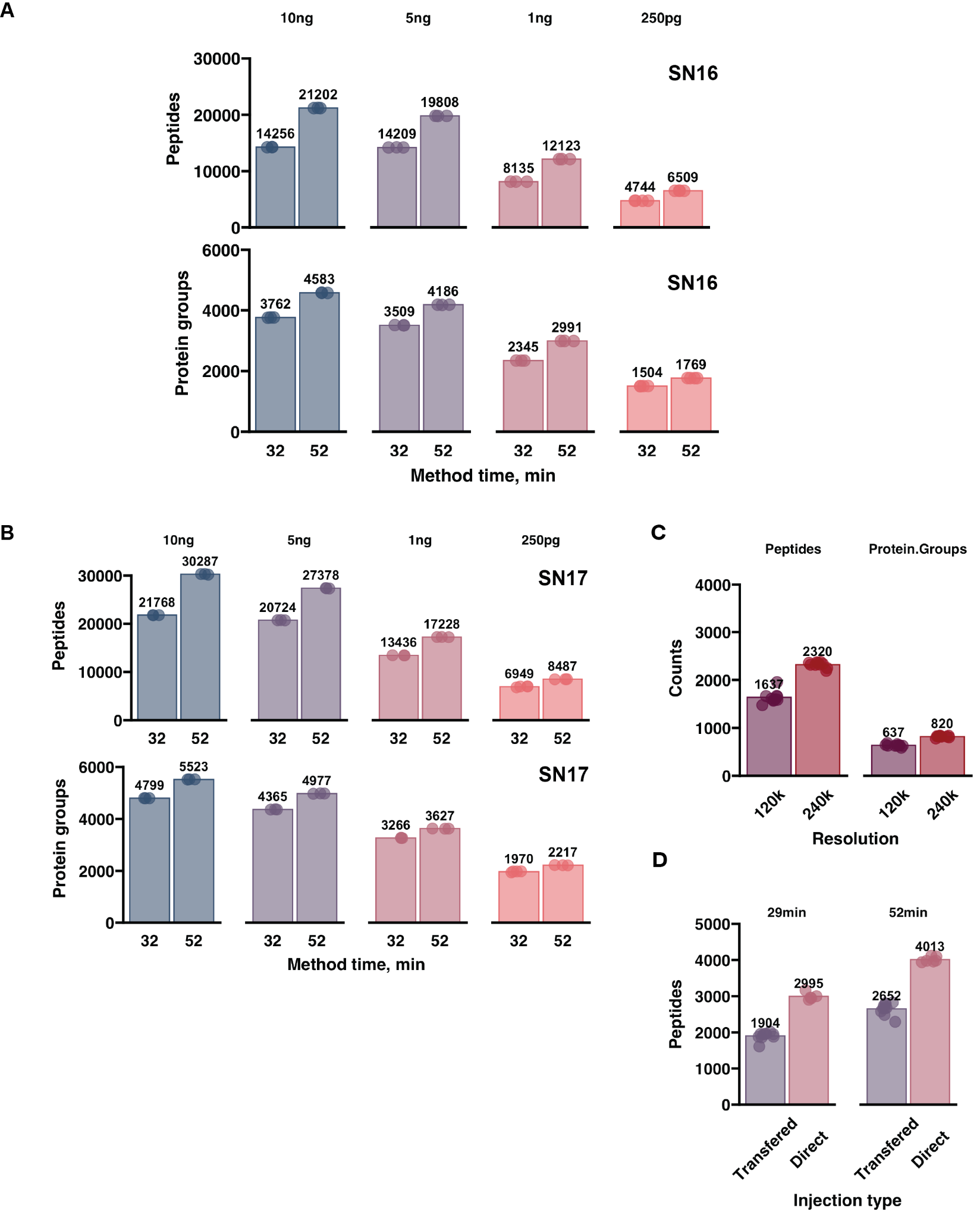

**Figure S4.** **uPAC column benchmarking supporting information. A)** Barplot showing detected number of proteins and peptides with the older Spectronaut16 version, with the pre-column configuration **B)** Barplots showing the detected number of proteins with the single-column and pre-column configurations, with Spectronaut16 “pre-release” version. **C)** Barplot showing the number of quantified peptides and protein groups with the 120k and 240k WISH-DIA methods. **D)** Barplot showing the number of quantified peptides from single-cell input with pre-column configuration. Related to Figure 4.

**Figure S5.** **DirectDIA detected protein accuracy with and without a library.** Density plots showing the log2 transformed light and heavy protein abundance ratios. Proteins quantified with directDIA shown in top and high-load (HL) library in the middle and gas-phase fractionated (GPF) in the bottom. Dashed lines denote expected ratios. MS1 based quantification used. Related to Figure 5.
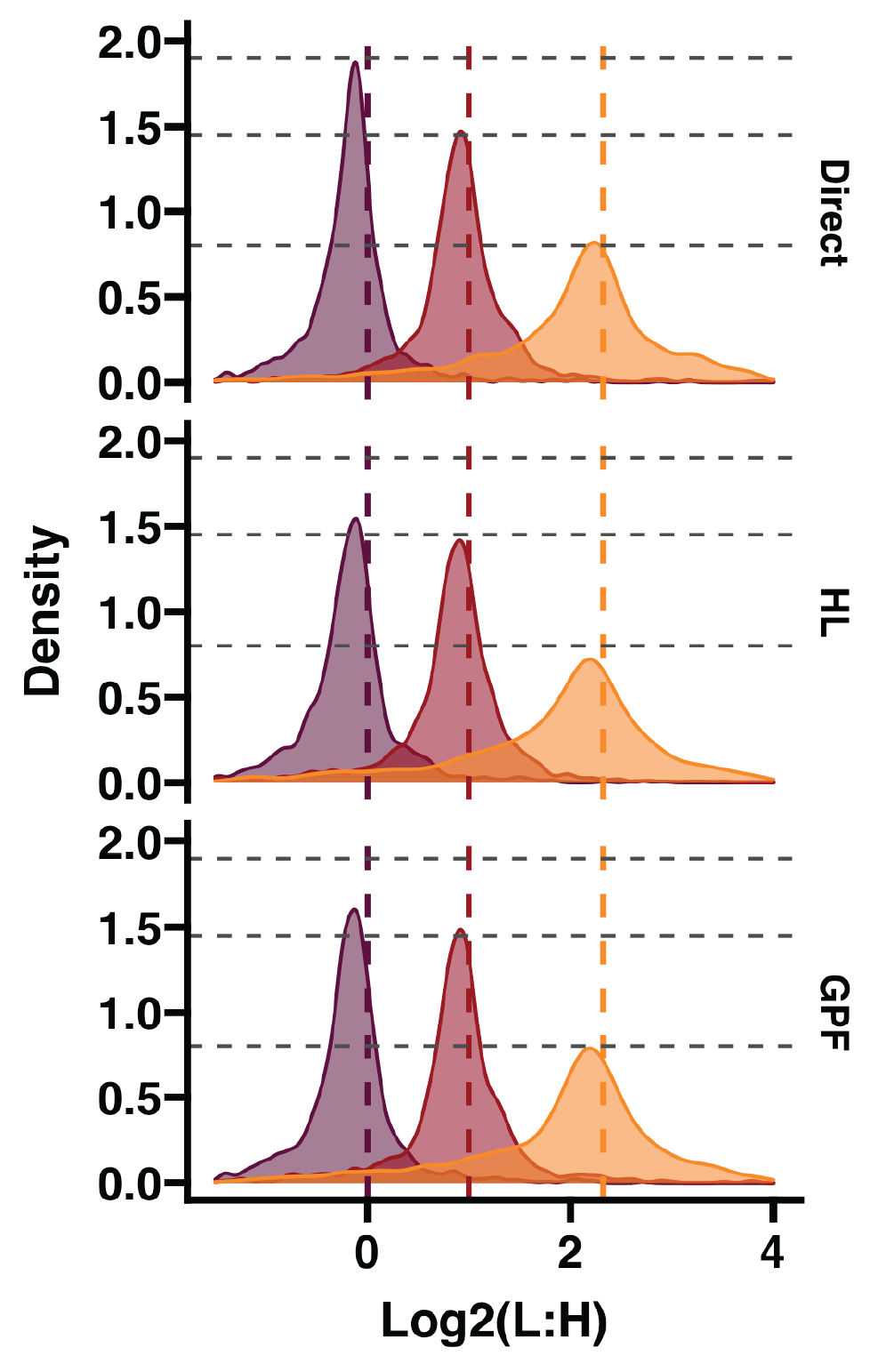

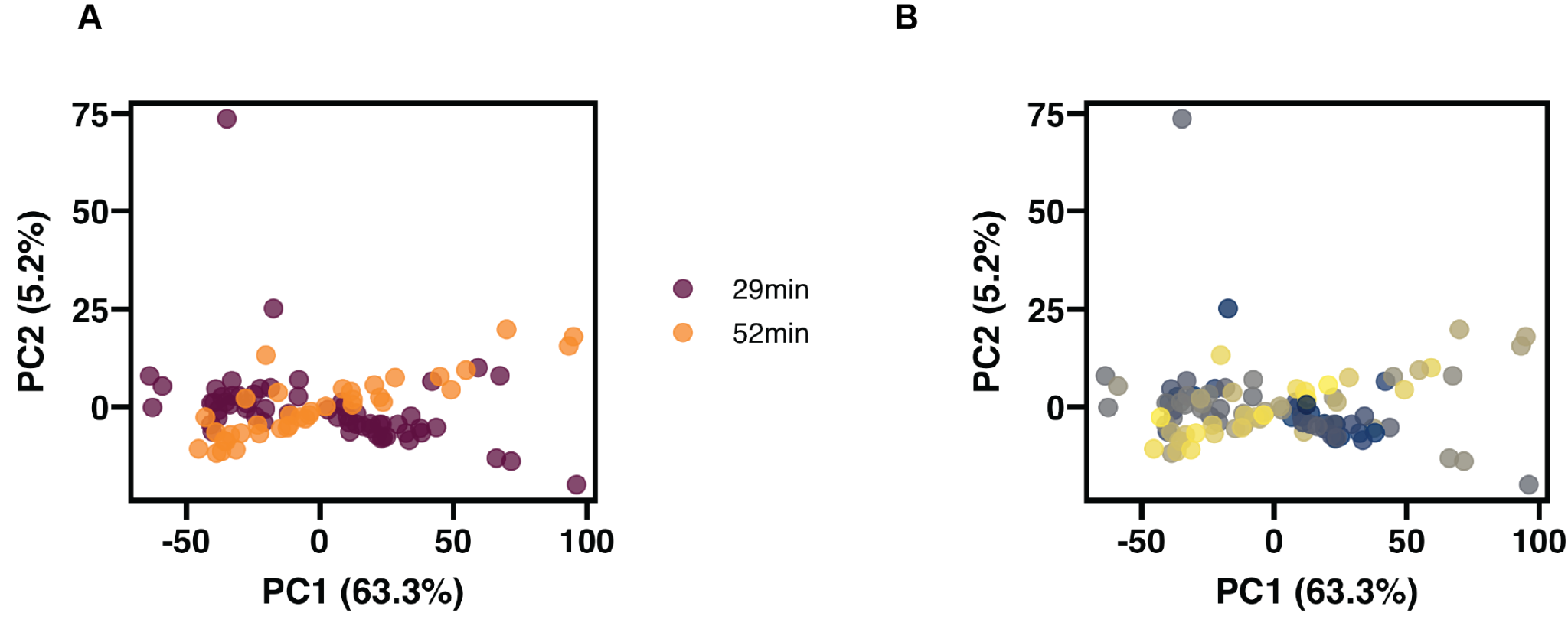

**Figure S6.** **Principal component analysis of single-cell datasets.** Clustering of the integrated single-cell with PCA. Principal components 1 and 2 are shown. **A)** Color coding marks the gradient used for the single-cell runs. **B)** Marks the order in which the samples were run. 52min runs were carried out after 29min.
